# Axl-mediated activation of TBK1 drives epithelial plasticity in pancreatic cancer

**DOI:** 10.1101/450049

**Authors:** Victoria H. Cruz, Emily N. Arner, Wenting Du, Alberto E. Bremauntz, Rolf A. Brekken

**Author notes:** These authors contributed equally to this work (Victoria H. Cruz, Emily N. Arner). Corresponding author: Rolf A. Brekken, PhD Hamon Center for Therapeutic Oncology Research University of Texas Southwestern Medical Center 6000 Harry Hines Blvd. Dallas, TX 75390-8593.

## Abstract

Pancreatic ductal adenocarcinoma (PDA) is characterized by an activating mutation in *KRAS*, which is critical for the biology of PDA progression. Direct inhibition of KRAS through pharmacological means remains a challenge; however, targeting key KRAS effectors has therapeutic potential. We investigated the contribution of TANK-binding kinase 1 (TBK1), a critical downstream effector of mutant active KRAS, to PDA progression. We report that higher levels of *TBK1* mRNA are associated with poorer overall survival in human PDA patients and that TBK1 supports the growth and metastasis of KRAS-mutant PDA by driving an epithelial plasticity program in tumor cells that enhances invasive and metastatic capacity. Further, we identify that the receptor tyrosine kinase Axl induces TBK1 activity in a Ras-RalB-dependent manner. These findings demonstrate that TBK1 is central to an Axl-driven epithelial-mesenchymal transition in KRAS-mutant PDA and suggest that interruption of the Axl-TBK1 signaling cascade above or below KRAS has potential therapeutic efficacy in this recalcitrant disease.

## Introduction

Pancreatic ductal adenocarcinoma (PDA) is a lethal and poorly understood human malignancy with patient survival that has not improved substantially in the last 40 years [1]. Each year, >45,000 new cases are diagnosed and an almost equal number of patients succumb to the disease [1]. The high incidence to death ratio is attributed to therapy resistance and to late diagnosis, at which point the tumor has metastasized [2].

Significant effort has gone into identifying critical pathways for PDA growth that can be exploited therapeutically. Understanding the biology of activating point mutations in the small GTPase, *KRAS*, which is an early genetic event in human PDA development and is present in 90% of PDA cases [3], has been a focal point of drug development strategies. Oncogenic KRAS is the dominant driver in PDA initiation and maintenance [4]; however, RAS itself has not been an amenable target for direct inhibition. As a result, developing therapeutic strategies that inhibit RAS effector signaling is attractive [5].

While the majority of RAS effector-targeted therapies are focused on the RAF and PI3K signaling networks, there is considerable evidence supporting the lesser studied, RALGEF-RAL effector pathway as a critical contributor to PDA growth [6-8]. Though not mutated as frequently as the RAF-MEK-ERK and PI3K signaling molecules in human cancer, the RALA and RALB GTPases of the RALGEF pathway are more consistently activated than RAF or PI3K in human pancreatic tumors [6-8]. The serine threonine protein kinase, TANK-binding kinase 1 (TBK1), is a major constituent of the RAL pathway and is critical to the development of Ras-driven cancers [9, 10]. Studies in lung cancer revealed that RALB activates TBK1, leading to the restriction of apoptosis, while having no effect on survival of non-tumorigenic epithelial cells [11]. Moreover, the expression of an oncogenic *KRAS* allele in TBK1-deficient murine embryonic fibroblasts was found to induce cell death, suggesting that TBK1 is integral for cells to tolerate transforming levels of oncogenic RAS [11]. The critical contribution of RALB and TBK1 to RAS-induced lung cancer growth was corroborated in an RNAi screen of synthetic lethal partners of oncogenic KRAS, where *RALB* and *TBK1* were identified as top hits [10].

Given the prominent activity of RalB in pancreatic cancer and the requirement of TBK1 for cells to tolerate transforming levels of oncogenic RAS, we hypothesized that TBK1 is critical to KRAS-driven pancreatic cancer growth. Here we show that TBK1 is expressed highly in pancreatic cancer and correlates with worse survival in human PDA patients. We found that the loss of TBK1 kinase function in preclinical mouse models of *KRAS*-mutant PDA resulted in reduced tumor load and reduced metastatic events, indicating that TBK1 activity contributes directly to the aggressive properties of pancreatic cancer. Taken together, our findings highlight TBK1 inhibition as a novel approach to targeting *KRAS*-mutant pancreatic cancer.

## Materials and methods

### Animals

NOD SCID mice were purchased from the UT Southwestern mouse breeding core (Dallas, TX). *Tbk1^Δ/Δ^, Tbk1*^+/+^, *Kras^LSL-G12D/+-^; Cdkn2a^Lox/Lox-^* (KI) and *Cdkn2a^Lox/Lox^*; *Ptf1a^Cre/+^* (IC) mice were generated as previously described [12-14]. *Tbk1^Δ/Δ^* and *Tbk1^+/+^* mice were used to breed with KI and IC mice to generate *Tbk1^+/+^*; *Kras^LSL-G12D/+^*; *Cdkn2a^Lox/Lox^*; *Ptf1a^Cre/+^* (*Tbk1^+/+^*: *KIC*) mice and *Tbk1^Δ/Δ^*: *KIC* mice. *LSL-Trp53^R172H/+^* mice were obtained from the National Cancer Institute Mouse Repository [15]. *Tbk1^Δ/Δ^* and *Tbk1^+/+^* mice were also used to breed with *Kras^LSL-G12D/+^*; *LSL-Trp53^LSL-R172H/+^* (KP) and *Ptf1a^Cre/+^* mice to generate *Kras^LSL-G12D/+^*; *LSL-Trp53^LSL-R172H/+^*; *Ptf1a^Cre/+^* (*KPC*) mice and *Tbk1^Δ/Δ^*: *KPC* mice. All mice were bred and maintained in a pathogen-free barrier facility with access to food and water *ad libitum*. All protocols for mouse use and euthanasia were reviewed and approved by the institutional animal care and use committee of the University of Texas Southwestern Medical Center (Dallas, TX).

### Animal studies

All experiments were conducted using littermate-controlled mice. All mice were fed a normal chow diet (16% protein diet, irradiated; Teklad Global Diets; Envigo, East Millstone, NJ). For endpoint studies, *Tbk1^+/+^*: *KIC* and *Tbk1*^Δ/Δ^: *KIC* mice were sacrificed and entire tissues, including pancreas/tumor, liver, lungs, and spleen, were harvested and weighed at 6, 8 and 10 weeks old, with n = 5-11 mice per time point per group. *Tbk1^+/+^*: *KPC* and *Tbk1*^Δ/Δ^: *KPC* mice were sacrificed between 4 and 5 months, with n ≥ 8 mice per time point per group. For all survival studies, mice were carefully monitored and sacrificed when they appeared moribund. For lung colonization studies, *Tbk1^+/+^*: *KIC* and *Tbk1*^Δ/Δ^: *KIC* cells (1 x 10^5^) were resuspended in 200 μl PBS and injected intravenously into the tail vein of 8-week-old female NOD SCID mice. Lungs were harvested at 7 or 12 days post injection and fixed in Bouin’s fixative for gross analysis of tumor nodules. Tumor colonization was analyzed by H&E.

### Histology and immunohistochemistry

Pancreas/tumors, livers, lungs, spleens, and kidneys were excised and fixed with 10% neutral buffered formalin solution overnight and embedded in paraffin for sectioning. All tissues were sectioned at 5 μm. After sectioning, slides were deparaffinized with xylene and rehydrated in decreasing ethanol dilution series and then stained with H&E. Masson’s Trichrome and Alcian Blue stains were performed on formalin-fixed, paraffin-embedded *KIC* tumor sections by the molecular pathology core (UT Southwestern, Dallas, TX). Sections for immunohistochemical analysis were blocked with 5% BSA and incubated with rabbit anti-CK19 (Abcam, ab15463) or rabbit anti-Vimentin (Cell Signaling, #5741) in blocking solution (5% BSA in TBS with 0.05% tween) at 4°C overnight. Horseradish peroxidase-conjugated donkey anti-rabbit IgG (Jackson ImmunoResearch Laboratories, West Grove, PA) was used as a secondary antibody for CK19 staining and TRITC-conjugated donkey anti-rabbit IgG (Jackson ImmunoResearch) was used as a secondary antibody for Vimentin staining. Negative controls included omission of primary antibody. All slides were visualized with a Nikon Eclipse E600 microscope (Nikon, Melville, NY) and color images were captured using a Nikon digital Dx1200me camera and ACT-1 software. Images were analyzed using NIS Elements AR 2.3 software (Nikon).

### RNA isolation and microarray analysis

Tumor tissues were excised from 8-week-old *Tbk1^Δ/Δ^* and *Tbk1^+/+^*: *KIC* mice and snap-frozen with liquid nitrogen (n = 3 tumors per genotype). Total RNA was isolated after tissue homogenization in TRIzol (Thermo Fisher, Waltham, MA) and RNA was extracted using an RNeasy RNA extraction kit (Qiagen, Germantown, MD). RNA was quantified using a NanoDrop instrument (Thermo Fisher) and checked for quality with a Bioanalyzer Instrument (Agilent). Gene expression was analyzed on a MouseWG-6 v2.0 Expression BeadChip (Illumina, San Diego, CA) through the UT Southwestern microarray core (Dallas, TX). Gene expression data analysis was performed through IPA software (Ingenuity Pathway Analysis, Qiagen). Java TreeView (Alok Saldanha) and Cluster 3.0 software (Michael Eisen, Berkeley Lab) were employed for hierarchical clustering gene expression analysis [16].

### Recombined *Cdkn2a* allele detection

Liver micrometastasis was assessed by quantitative RT-PCR (qPCR) for the recombined *Cdkn2a*(*Ink4a*/*Arf*) allele. Briefly, frozen livers were homogenized in SDS lysis buffer (100 mM Tris pH 8.8, 5 mM EDTA, 0.2% SDS, 100 mM NaCl) and digested at 56¼C overnight. DNA was extracted using phenol-chloroform-isoamyl-alcohol (25:24:1) and quantitative RT-PCR was performed using iQ SYBR Green Supermix (Bio-Rad, California, USA). The following validated primers were used for analysis of CDKN2A: gccgacatctctctgacctc (forward) and ctcgaaccaggtttccattg (reverse). Each sample was analyzed in triplicate.

### Immunoblotting

Tissues and cells were lysed in ice-cold RIPA buffer (50 mM Tris-Cl, 150 mM NaCl, 1% Nonidet P-40, 0.5% sodium deoxycholate, and 0.1% SDS) containing cocktails of protease (Thermo Fisher) and phosphatase inhibitors (Sigma-Aldrich, St. Louis, MO) and centrifuged for 20 min at 13,000 × *g* at 4°C. Total protein concentration was calculated using a bicinchoninic acid assay kit (Thermo Fisher). Proteins were resolved by SDS-PAGE and transferred to a methanol-activated polyvinylidene difluoride membrane. All primary and secondary antibodies were diluted in 5% donkey serum in TBS with 0.05% tween. Primary antibodies used included the following: anti-pAKT(S473; Cell Signaling, #4060), anti-AKT (Cell Signaling, #9272), anti-pAxl (Y779; R&D, AF2228), anti-Axl (Santa Cruz, sc-1096), anti-Claudin1 (Cell Signaling, #13255), anti-E-cadherin (Cell Signaling, #3195), anti-IKKε (Biochain, Z5020108), anti-IRF3 (Santa Cruz, sc-9082), anti-pIRF3 (S396; Cell Signaling, #4947), anti-N-cadherin (Cell Signaling, #13116), anti-P65 (Santa Cruz, #8008), anti-pP65 (Cell Signaling, #3033) anti-Ras (Abcam, ab108602), anti-Snail (Cell Signaling, #3879), anti-Slug (Cell Signaling, #9585), anti-pTBK1 (S172; Cell Signaling, #5483 and Abcam, #109272), anti-TBK1 (Abcam, ab40676), anti-Vimentin (Cell Signaling, #5741), anti-Zeb1 (Cell Signaling, #3396), anti-ZO-1 (Cell Signaling, #8193). Anti-β-actin (Sigma-Aldrich, A2066), anti-GAPDH (Cell Signaling, #2118) and anti-Tubulin (Abcam, ab4047) were used as loading controls for all Western blots shown. Horseradish peroxidase-conjugated donkey anti-rabbit, donkey anti-mouse, and donkey anti-goat IgG (1:10,000; Jackson ImmunoResearch) were used as a secondary antibodies. Membranes were exposed with Clarity Western ECL Blotting Substrate (Bio-Rad) and visualized with the Odyssey Fc imager (LI-COR Biotechnology, Lincoln, NE).

### Cell lines

Human and mouse cancer cell lines (AsPC-1, Capan-1, Hs766T, MCF7, MIA PaCa-2, PANC-1, PL-45) were obtained from ATCC (Manassas, VA). HPNE (human pancreatic nestin-expressing) cells were generated as previously described [17] and obtained from the UT MD Anderson Cancer Center (Houston, TX). The *KPC*-M09 and *KPfC-8* cell lines were isolated from spontaneous tumors originating in a *KPC* and *KPfC* (*Kras^LSL-G12D/+^; Trp53^lox/lox^; Ptf1a^Cre/+^*) mouse, respectively, as previously described [18]. All cell lines were cultured in DMEM or RPMI (Invitrogen, Carlsbad, CA) containing 10% FBS and maintained in a humidified incubator with 5% CO2 at 37°C. The human cell lines were DNA fingerprinted for provenance using the Power-Plex 1.2 kit (Promega) and confirmed to be the same as the DNA fingerprint library maintained by ATCC. All cell lines were confirmed to be free of *mycoplasma* (e-Myco kit, Boca Scientific) before use.

Isogenic cell lines were derived from individual tumors of 8-week-old *Tbk1^+/+^: KIC* and *Tbk1^Δ/Δ^: KIC* mice. Each tumor was minced and digested with 1 % collagenase type I, DMEM, 10 mM HEPES, and 1% FBS at 37°C to obtain a single-cell suspension. Cell suspensions were centrifuged at low speed to pellet large debris, resuspended in wash buffer, and passed through a 70 μm cell strainer. The resulting cell suspension was plated at low density to isolate tumor cell populations using cloning rings. Cells were confirmed to be tumor cells by immunocytochemistry and PCR. These cell lines were expanded and stained for tumor cell markers. Cell lines were confirmed to be pathogen-free before use. Clones *Tbk1^+/+^: KIC-A, Tbk1^+/+^: KIC-B, Tbk1^+/+^: KIC-D, Tbk1^Δ/Δ^: KIC-A, Tbk1^Δ/Δ^: KIC-B*, and *Tbk1^Δ/Δ^: KIC-C* were used in subsequent experiments. Cells were cultured in DMEM containing 10% FBS and maintained at 37°C in a humidified incubator with 5% CO2 and 95% air.

### *In vitro* drug response assay

Assays were performed in 96-well format as described [19]. Briefly, cells were plated on day 0 and compound II [9] was added on day 1 in 4-fold dilutions starting at 20 µM (highest dose). For each assay, 8 different drug concentrations were tested with 8 replicates per concentration. Relative cell number was determined by adding MTS (Promega, Madison, WI, final concentration 333 µg/ml), incubating for 1 to 3 hours at 37°C, and reading absorbance at 490 nm on a plate reader (Spectra Max 190, Molecular Devices, Downingtown, PA). Drug sensitivity curves and IC50s were calculated using in-house software. Response was validated in replicate plates (n ≥ 4).

### Wound healing and invasion assays

Wound healing assays were conducted in 6-well plates. Monolayers of cells were grown in low-serum media until 90% confluency was reached. Each well was scratched with a P200 pipette tip to create an artificial wound, washed with PBS to remove residual cells and replaced with fresh media containing 10% FBS. Cells were photographed at indicated time points after wounding. Wound closure was measured as a percentage of original wound width with MRI Wound Healing Tool macro (ImageJ).

Invasion assays were carried out with QCM ECMatrix Cell Invasion Assays (EMD Millipore, Burlington, MA). In brief, cells were serum-starved overnight and then seeded the next day on transwell inserts (8 μm pore size) that were lined with a reconstituted basement membrane matrix of proteins derived from the Engelbreth Holm-Swarm mouse tumor. The inner chambers were filled with serum-free medium while the outer chambers were filled with medium containing 10% FBS as the chemoattractant. After the indicated time points, invaded cells on the bottom of the insert membrane were dissociated from the membrane when incubated with cell detachment buffer and subsequently lysed and detected by CyQuant GR dye.

### Organotypic culture and immunocytochemistry

For each cell line, 2000 cells were plated in 8-well chamber slides onto a base layer of Matrigel (5 mg/ml) and collagen I (1.5–2.1 mg/ml) and cultured for 3 to 4 days in a humidified 37°C incubator as previously described [20]. For immunocytochemistry, cultures were fixed in 2% formalin (Sigma-Aldrich) in PBS for 20 min, permeabilized with 0.5% Triton X-100 in PBS for 10 min at room temperature, incubated with Alexa Fluor 488 Phalloidin (A12379, Invitrogen) or Alexa Fluor 546 Phalloidin (A22283, Invitrogen) in immunofluorescence buffer as described [21] for 1 hour at room temperature and mounted using ProLong Gold antifade reagent with DAPI (Invitrogen). Images were acquired using a confocal laser-scanning microscope (LSM880, Zeiss) through UT Southwestern’s live cell imaging core (Dallas, TX) and a Nikon Eclipse E600 microscope with a Nikon Digital Dx1200me camera.

### Active GTPase assays

Active RAS in cell lysates was measured via precipitation with GST-tagged RAF-RBD beads (Ras Pull-down Activation Assay Biochem Kit; Cytoskeleton, Denver, CO). Active RalB in cell lysates was measured via precipitation with RalBP1 PBD Agarose beads (Cell BioLabs, San Diego, CA). Lysates were prepared and precipitation was performed per manufacturer instructions. Subsequently, pull-down samples and respective whole cell lysates were immunoblotted with anti-RAS (pan) or anti-RALB (pan) and indicated loading controls.

### Reagents

The mammalian expression plasmid pCDH-CMV-MCS-TBK1-EF1-NEO was generously provided by Drs. Peiqing Shi and James Chen (UT Southwestern Medical Center, Dallas, TX). Lentiviral-based expression constructs were packaged by co-transfection of HEK293T cells with psPAX2 and pMD2.G packaging system (4:2:1). Polyethylenimine (PEI) was used for transfection at a 3:1 ratio of total DNA. Transfection media was replaced with 10% complete DMEM 24 hours post transfection, and incubated a further 24 hours prior to viral particle collection. *Tbk1^+/+^* and *Tbk1^Δ/Δ^: KIC* cells were seeded at a density of 1 x 10^6^ cells per 10 cm dish. Twenty-four hours later, the cells were infected with lentiviral particles and polybrene (10 μg/ml). At 24 hours post infection, cells were given fresh medium containing G418 (400 μg/mL, InvivoGen) for selection and maintained in culture under selection for 3 weeks following initial infection. AF854 (R&D Systems, Minneapolis, MN) was previously shown to activate mouse Axl and was used at indicated concentrations to stimulate mouse Axl [22]. Compound II was synthesized by William G. Bornmann (UT MD Anderson Cancer Center) and found to be a potent inhibitor of TBK1 and IKKε in a chemical compound screen as previously described [9].

### Clinical data set

Gene expression data and survival analyses of 84 pancreatic cancer patients with annotated clinical outcomes were downloaded from the Cancer Genome Atlas. For survival analyses, gene expression data was divided into the top 25% and bottom 75%. Cox regression was used to calculate hazard ratios and Kaplan-Meier survival analyses. Significance for these plots was determined using the log-rank test.

### Statistics

Statistical analyses were performed using GraphPad Prism (GraphPad, La Jolla, CA). Results are expressed as mean ± SEM. All data were analyzed by *t* test. Significance was accepted at *p* < 0.05, with asterisks denoting *p*-value levels: \**p* < 0.05, \*\**p* < 0.01, \*\*\**p* < 0.001, and \*\*\*\**p* < 0.0001.

## Results

### TBK1 expression in pancreatic cancer

TBK1 is expressed in numerous epithelial tumors, including breast, lung, and colon [10, 23-25]. However, the function and activity of TBK1 in human pancreatic cancer has not been characterized extensively. We found that TBK1 was expressed and was active (pTBK1) under basal conditions in a panel of human *KRAS*-mutant PDA cell lines. In contrast, TBK1 was expressed but with reduced activity in normal pancreatic ductal epithelial cells (HPNE) [17] or *KRAS wild-type* PDA cells. Mouse embryonic fibroblasts isolated from *Tbk1*-mutant mice (*Tbk1^Δ/Δ^*) [12] served as a negative control with no detectable expression of TBK1 or pTBK1 **(Fig. 1a)**. Additionally, we observed higher TBK1 protein levels in spontaneous pancreatic tumors from genetically engineered mouse models (GEMMs) compared to normal pancreas from littermate controls **(Fig. 1b)**. We previously reported that *TBK1* expression is associated with a poor prognosis in pancreatic cancer patients (stages I-II) from the Cancer Genome Atlas [26]. From this cohort (n = 84), gene expression was divided into top 25% and bottom 75% percentiles for Kaplan-Meier survival analyses. While expression of TBK1 homolog, *IKBKE* showed no correlation with survival, high expression of *TBK1* showed a strong trend towards poorer overall survival in this patient cohort (*p* = 0.07) **(Fig. 1c-d)**. Though not causative, this data suggests that TBK1 expression and activity is important in human pancreatic cancer.

**Figure 1.**
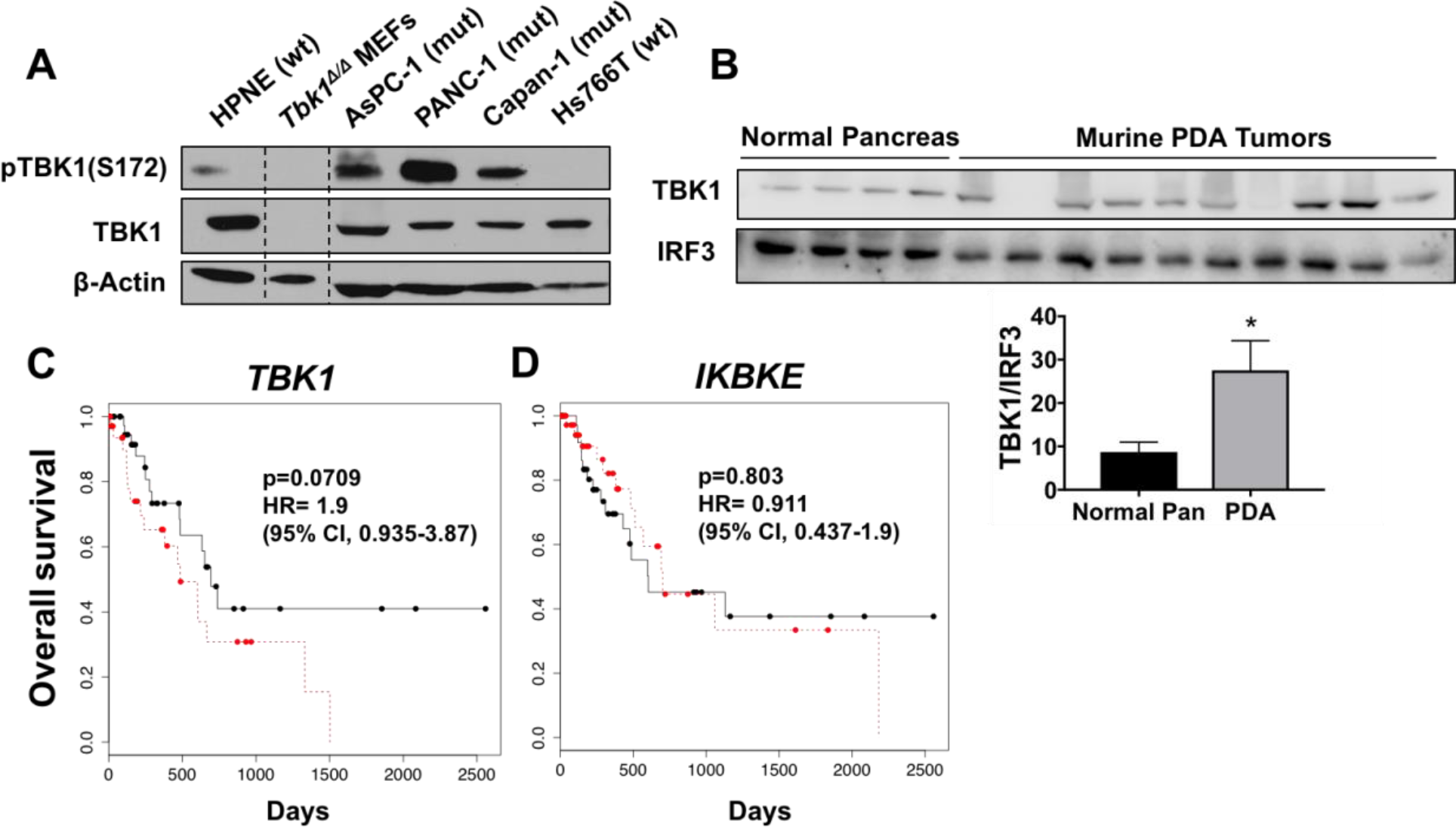
TBK1 is highly expressed in pancreatic cancer and correlates with worse overall survival. **a)** Total and pTBK1 (S172) expression in wild-type *KRAS* (wt) or mutant active *KRAS* (mut) human normal (HPNE) or pancreatic ductal adenocarcinoma (PDA) cell lines. *Tbk1^Δ/Δ^* mouse embryonic fibroblasts (MEFs) served as a negative control. β-actin was used as a loading control. The dotted line indicates where blot was cropped; however, all samples were run on the same gel and exposed simultaneously. **b)** Total TBK1 expression in murine PDA tumors relative to normal pancreas from nontumor-bearing littermate controls. Total IRF3 was used as a loading control. Intensity quantification is representative of mean +/- SEM. Unpaired *t* test, \**p* < 0.05. *TBK1* (**c**) and *IKBKE* (**d**) gene expression (mRNA) in pancreatic cancer patient survival data from the Cancer Genome Atlas (25). Gene expression was divided into top 25% (red) and bottom 75% (black) percentiles. For statistical analysis, Cox regression was used to calculate hazard ratios and Kaplan-Meier survival analyses.

### KRAS-driven pancreatic cancer growth is disrupted by restricting TBK1 kinase activity

After observing high TBK1 activity in human PDA cell lines with mutant active *KRAS*, we tested cell line dependency on TBK1 for viability using compound II, a 6-aminopyrazolopyrimidine derivative with an IC50 of 13 nM and 59 nM against TBK1 and IKKε, respectively [9]. Compound II has 100- to 1000-fold less activity against other purified recombinant protein kinases tested, including PI3K and mTOR family members [9]. Following a 96-hour exposure to compound II, the viability of human and mouse PDA cell lines was measured using a MTS assay **(Fig. 2a)**. Cell lines harboring a *KRAS* mutation (shown in red) segregated distinctly from wild-type *KRAS* cell lines (shown in blue) with an IC50 of ∼1 μM versus greater than 5 μM, respectively. The *KRAS* wild-type PDA cell line, Hs766T, which previously showed no TBK1 activity (**Fig. 1a**), was the least sensitive to TBK1 inhibition with an IC50 above 20 μM. In this panel, MCF-7 is a *KRAS* wild-type cancer cell line that was previously shown [9] to be less sensitive to compound II and therefore served as a negative control with an IC_50_ of 5 μM. Additionally we tested two murine cell lines (*KPC*-M09 and *Tbk1^+/+^: KIC-B*) isolated from mutant Kras-driven PDA GEMMs that proved to be more sensitive to compound II than the human PDA cell lines. *Tbk1^Δ/Δ^: KIC-A* cells were isolated from a PDA GEMM that lacks functional TBK1 and therefore served as an additional negative control with an IC50 of 7 μM. The selective sensitivity of mutant *KRAS* PDA cell lines to TBK1 inhibition in culture led us to speculate whether TBK1 was essential to RAS-driven PDA tumorigenesis in vivo.

**Figure 2.**
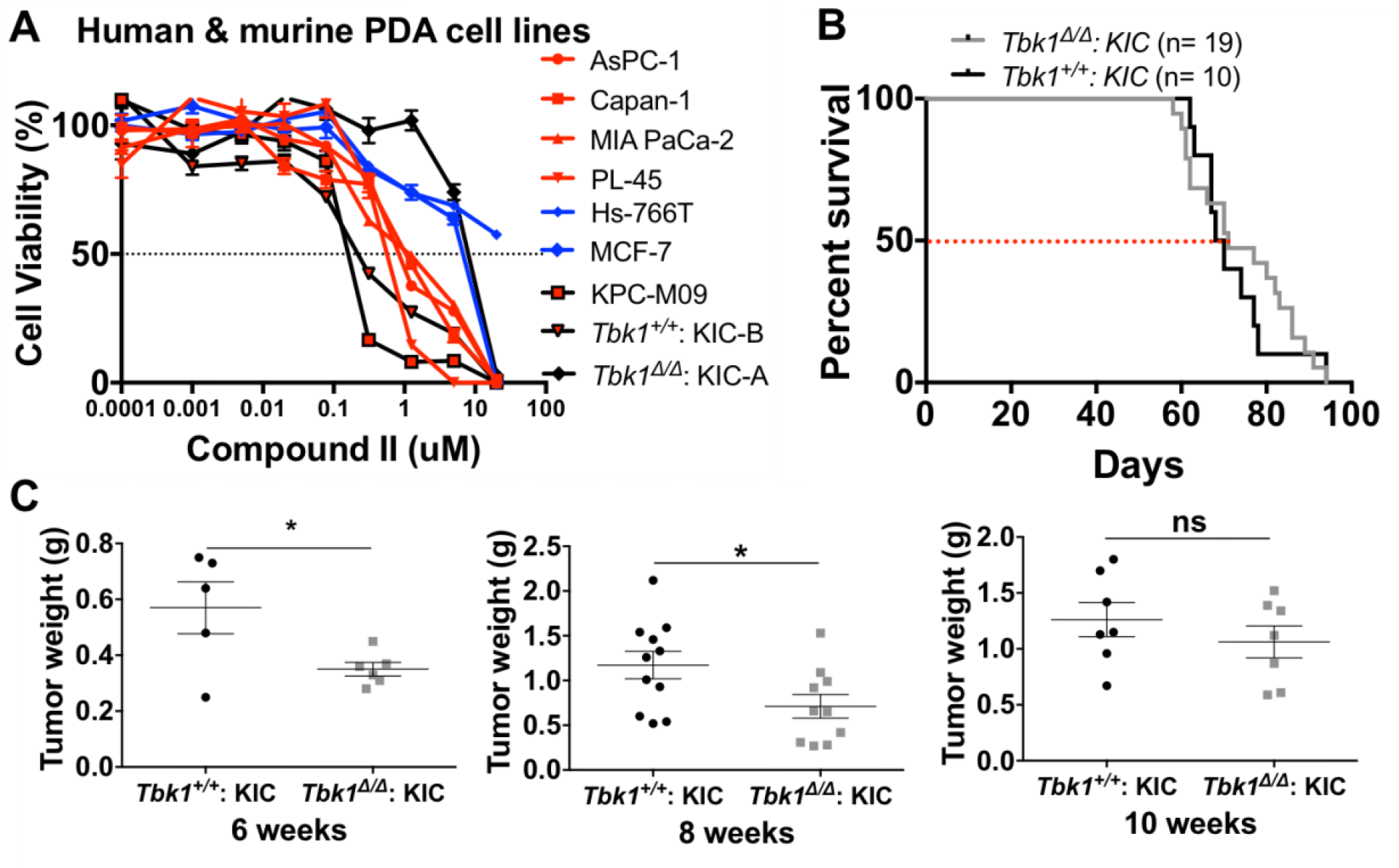
TBK1 promotes pancreatic ductal adenocarcinoma (PDA) tumor growth. **a)** Dose response curves of human and murine PDA cell lines treated with the TBK1 inhibitor compound II. Red indicates a human *KRAS* -mutant cell line, blue indicates a human *KRAS* wild-type cell line, and black indicates a murine PDA cell line. Error bars represent SEM from triplicate experiments. **b)** Kaplan-Meier survival curve of *Tbk1^Δ/Δ^*: *KIC* and *Tbk1^+/+^*: *KIC* mice. A log-rank Mantel-Cox test was used for survival comparison. **c)** Endpoint tumor weights at 6, 8, and 10 weeks in *Tbk1^Δ/Δ^*: *KIC* and *Tbk1^+/+^*: *KIC* mice. **d)** Metastasis to the liver was quantified by qPCR for recombined *Cdkn2a* allele (n = 5-6 mice/group). Results are representative of mean +/- SEM. Unpaired *t* test, \**p* < 0.05; *ns*, not significant.

To assess the contribution of TBK1 to PDA progression in vivo, we crossed *Tbk1* mutant mice *(Tbk1Δ/Δ)* that harbor two copies of a null *Tbk1* allele into a GEMM of PDA. The “null” *Tbk1* allele encodes a truncated TBK1 protein that is kinase-inactive and expressed at low levels, thereby allowing analysis of global TBK1 kinase loss in vivo [12]. *Tbk1^Δ/Δ^* animals were crossed with *KIC* (*Kras^LSL-G12D/+^*; *Cdkn2a^Lox/Lox^*; *Ptf1a^Cre/+^*) mice. *KIC* mice present with low-grade ductal lesions by 3 weeks of age [13, 14, 27] that progress to pancreatic adenocarcinomas such that all mice are moribund between 7 and 11 weeks of age. We hypothesized that TBK1 was critical for RAS-mediated oncogenesis in pancreatic cancer, thus the expectation was that *Tbk1^Δ/Δ^*: *KIC* mice would have smaller tumors and outlive *Tbk1^+/+^*: *KIC* mice. In comparing tumor sizes, we observed that tumors from *Tbk1^Δ/Δ^*: *KIC* mice were between 20%-40% smaller than *Tbk1^+/+^*: *KIC* tumors at multiple time points, yet there was no difference in overall survival between the two groups **(Fig. 2b-c, Supplemental Fig. 1a)**. Malnutrition resulting from loss of normal exocrine pancreas function and pancreatic enzyme insufficiency contributes to early death in the *KIC* model [28]. Thus, we cannot exclude the possibility that the antitumor effects of *Tbk1* loss are surpassed by the aggressiveness of the *KIC* model, causing the mice to succumb to malnutrition.

### *Tbk1^Δ/Δ^: KIC* tumors are differentiated

To better understand TBK1-dependent mechanisms of tumor cell growth contributing to larger tumors in *KIC* mice, we performed gene expression analysis using RNA isolated from 8-week-old *Tbk1^Δ/Δ^*: *KIC* and *Tbk1^+/+^*: *KIC* tumors. One of the most significant and top dysregulated gene networks between *Tbk1^Δ/Δ^*: *KIC* and *Tbk1^+/+^*: *KIC* tumors identified by Ingenuity Pathway Analysis (Qiagen) was the “cancer/cellular movement” network. This network included a large number of genes involved in epithelial-to-mesenchymal transition (EMT). In comparison to *Tbk1^+/+^*: *KIC* tumors, all three *Tbk1^Δ/Δ^*: *KIC* tumors showed a trend of lower expression of mesenchymal genes, such as vimentin and matrix metallopeptidase 9 (MMP9), and higher expression of epithelial genes including claudin 3, 4, and 10 and tissue inhibitor of metalloproteinase 3 (TIMP3) **(Fig. 3a)**. Low vimentin expression was confirmed at the protein level by immunofluorescent staining in tumors from 8-week-old animals **(Fig. 3b)**. Further, this EMT gene expression signature was consistent with alcian blue staining of *Tbk1^Δ/Δ^*: *KIC* and *Tbk1^+/+^*: *KIC* tumors. Alcian blue stain mucins that are expressed by ductal epithelial cells in panIN (pancreatic intraepithelial neoplasia) lesions [27]. Representative images show that *Tbk1^Δ/Δ^*: *KIC* tumors contain more alcian blue positive epithelial cells compared to less differentiated *Tbk1^+/+^*: *KIC* tumors **(Fig. 3c)**. Furthermore, we evaluated collagen deposition, a hallmark of pancreatic cancer that promotes EMT in PDA and is upregulated in response to epithelial plasticity [27, 29]. *Tbk1^+/+^*: *KIC* tumors showed higher levels of fibrillar collagen than *Tbk1^Δ/Δ^*: *KIC* tumors by trichrome histology **(Fig. 3d)**.

**Figure 3.**
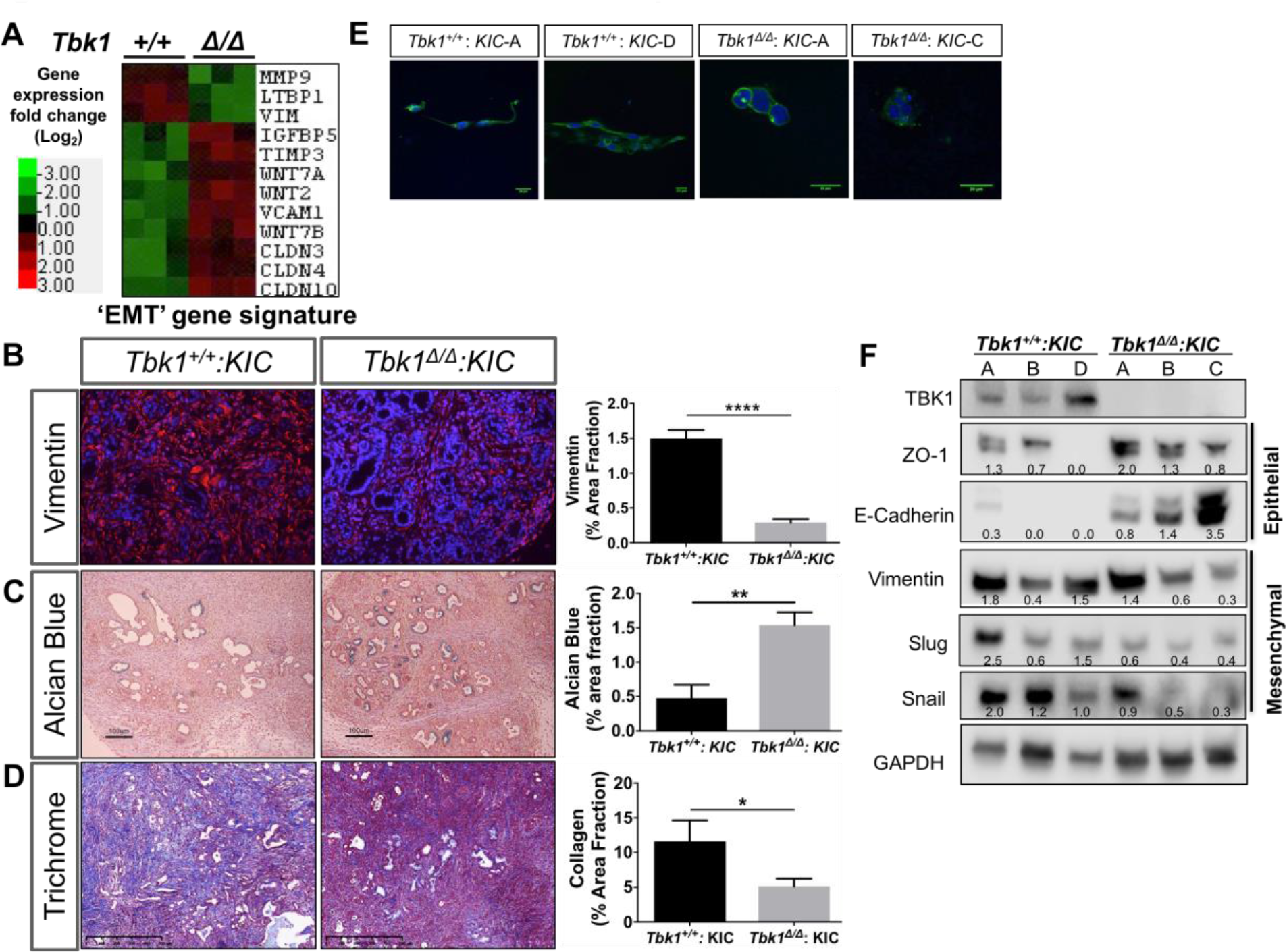
*Tbk1 ^Δ/Δ^*: *KIC* tumors are more “epithelial.” **a)** Heatmap representing gene expression fold change (log2) of epithelial-to-mesenchymal transition (EMT)-related genes from *Tbk1^+/+^: KIC* and *Tbk1^Δ/Δ^: KIC* tumors. Color key indicates gene expression fold change; n = 3 tumors/genotype, *p* < 0.05 for all genes between *Tbk1^+/+^: KIC* and *Tbk1^Δ/Δ^: KIC* tumors. **b-d)** Representative images of vimentin **(b)**, alcian blue **(c)**, and trichrome **(d)**-stained tumors from 8-week-old *Tbk1^+/+^: KIC* and *Tbk1^Δ/Δ^: KIC* mice. Scale bars indicate 100 μm for alcian blue images and 500 μm for trichrome images; n ≥ 4 mice/group, 4 images/mouse. Results are representative of mean +/- SEM. Unpaired *t* test, \**p* < 0.05, \*\**p* < 0.01, \*\*\*\**p* < 0.0001. **e)** Representative confocal images of single-cell clones isolated from *Tbk1^+/+^: KIC* and *Tbk1^Δ/Δ^: KIC* tumors plated on a mixed layer of collagen and Matrigel. Nuclei are labeled with DAPI (blue) and F-actin is labeled with phalloidin (green). Scale bars indicate 20 μm. **f)** Protein lysates isolated from *Tbk1^+/+^: KIC* and *Tbk1^Δ/Δ^: KIC* cell lines were immunoblotted for indicated epithelial and mesenchymal markers. GAPDH was used as a loading control.

To confirm the tumor cell epithelial phenotype observed in *Tbk1^Δ/Δ^*: *KIC* tumors, we isolated single cell clones from *Tbk1^+/+^*: *KIC* and *Tbk1^Δ/Δ^*: *KIC* tumors. In total, 3 cell lines per genotype were generated, each from individual tumors. In accordance with gene expression data from *Tbk1^+/+^*: *KIC* tumors, each cell line isolated showed evidence of EMT with an elongated spindle-like cell shape, a characteristic often associated with mesenchymal cells. Moreover, cell lines from *Tbk1^Δ/Δ^*: *KIC* tumors exhibited a “cobblestone” morphology, a feature consistent with epithelial cells. These differences in morphology were observed in organotypic culture after the cells were plated on a mixed layer of collagen and Matrigel **(Fig. 3e)**. Evaluation of EMT-related markers revealed higher expression of epithelial proteins, ZO-1 and E-cadherin and lower expression of the mesenchymal proteins Vimentin, Slug and Snail in *Tbk1^Δ/Δ^*: *KIC* cell lines **(Fig. 3f)**. These results illustrate a unique epithelial signature in *Tbk1^Δ/Δ^*: *KIC* cells and have implications for functional differences in tumor cell motility.

### *Tbk1^Δ/Δ^: KIC* tumor cells are less migratory and invasive

Epithelial plasticity changes commonly correspond with alterations in tumor cell motility and invasiveness [30]. Given that *Tbk1^Δ/Δ^*: *KIC* tumors and cell lines are less mesenchymal in gene expression and morphology, we hypothesized that functional TBK1 is important for tumor cell migration. To compare motility and invasiveness between *Tbk1^+/+^*: *KIC* and *Tbk1^Δ/Δ^*: *KIC* tumor cell lines, we performed a series of wound healing and transwell invasion assays. Despite the fact that *Tbk1^Δ/Δ^*: *KIC* cell lines proliferate more quickly in culture **(Supplemental Fig. 2b)**, they did not migrate as well as *Tbk1^+/+^*: *KIC* cells **(Fig. 4a)**. For each cell line, migration at 24 hours was plotted against total cell population doublings **(Fig. 4b)**. The negative correlation in this plot demonstrates that enhanced migration seen in *Tbk1^+/+^*: *KIC* cells is not a result of faster growth rates. ECM-coated transwell migration assays also revealed a 20%-50% decrease in invasive capacity in *Tbk1^Δ/Δ^*: *KIC* cell lines when compared to *Tbk1^+/+^*: *KIC* cells at multiple time points (24 and 72 hours) **(Fig. 4c-d)**. These results further highlight the reduced migratory ability of *Tbk1^Δ/Δ^*: *KIC* cells.

**Figure 4.**
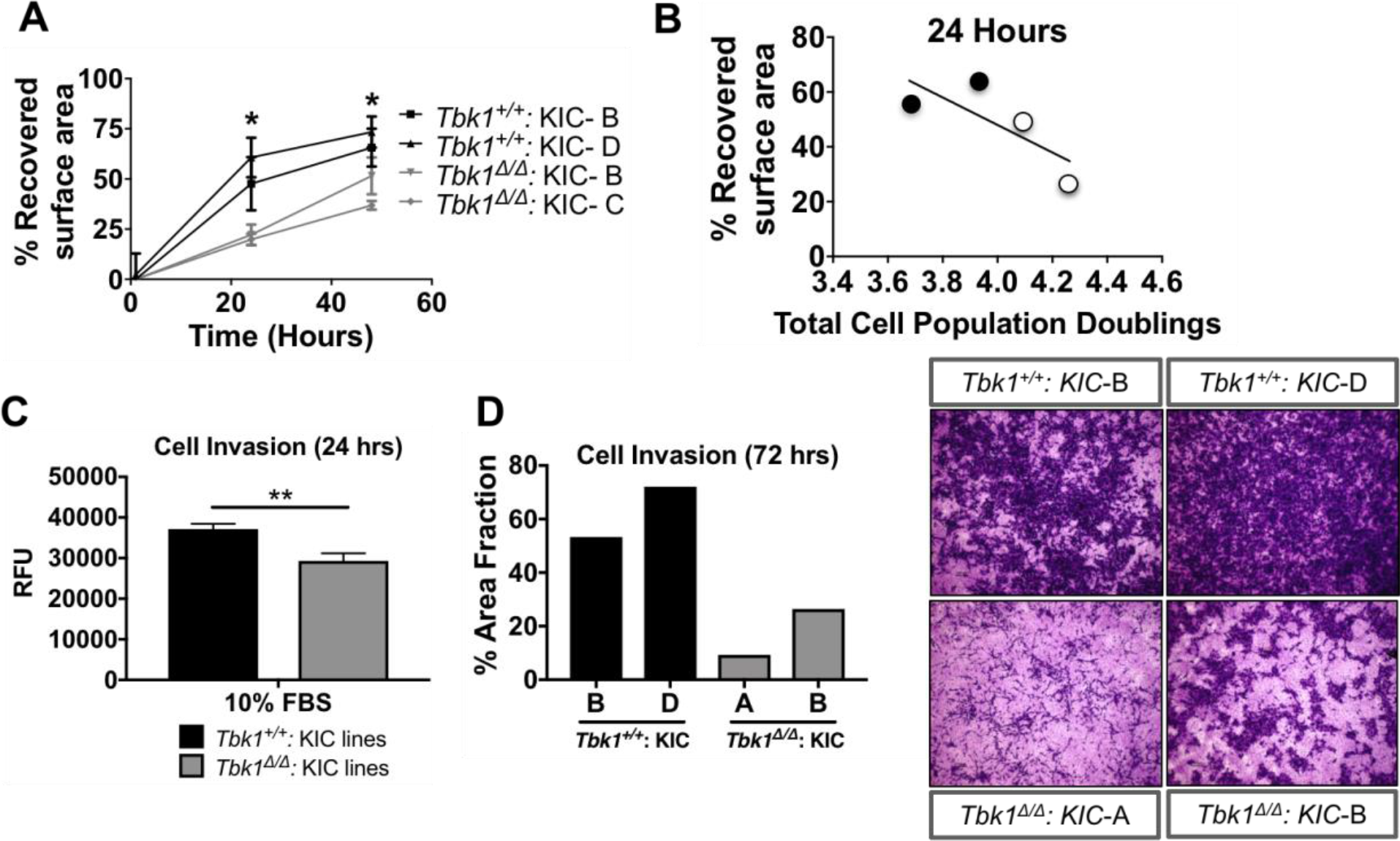
*Tbk1^Δ/Δ^: KIC* tumor cells are less migratory and invasive than *Tbk1^+/+^*: *KIC* cells. The migratory capacity of cell lines derived from *Tbk1: KIC* tumors was investigated using scratch **(a, b)** and transwell migration **(c, d)** assays. **a)** Recovered surface area of *Tbk1*: *KIC* cell lines at 0, 24 and 48 hours post scratch in wound healing assay. Quantification is representative of 3 wells/cell line. **b)** Total cell population doublings (from initial plating) were calculated for each cell line in **(a)** and correlated to migration. Black dots indicate *Tbk1^+/+^: KIC* cell lines and grey dots indicate *Tbk1^Δ/Δ^: KIC* tumor cells. **c)** Quantification (expressed as relative fluorescent units) of *Tbk1*: *KIC* cells after migration through ECM-coated transwell membranes 24 hours after plating in serum-free media. **d)** Quantification and images of crystal violet-stained *Tbk1*: *KIC* cell lines 72 hours post plating in the transwell assay. Quantification was measured as a percent area fraction of membrane covered in cells (stained in purple). **c, d)** The lower chamber of the transwell were filled with 10% FBS containing media as the chemoattractant. Results are representative of mean +/- SEM. Unpaired *t* test, \**p* < 0.05, \*\**p* < 0.01.

### Evaluation of *Tbk1* loss on pancreatic cancer metastases

Next we asked whether the reduction in tumor cell motility with kinase-dead *Tbk1* translated to fewer metastases in vivo. The *KIC* mouse model of PDA is an aggressive model with an average life span of ∼10 weeks [13, 14, 27]. As such, these mice rarely develop gross metastases, making this model less than ideal for comparing metastatic burden. However, it’s worth mentioning that livers from *Tbk1^+/+^* (n = 14) and *Tbk1^Δ/Δ^* (n = 12) *KIC* mice were examined for micro-metastases using histology and qPCR. Lesions were identified in 6 livers from *Tbk1^+/+^* animals but no lesions were found in livers from *Tbk1*-mutant mice **(Supplemental Fig. 1b-d)**. To more robustly study the effect of *Tbk1* loss on metastatic potential, we employed two different animal models. First, we exploited an experimental metastasis model where *Tbk1^+/+^* or *Tbk1^Δ/Δ^*: *KIC* cell lines were injected intravenously (i.v.) into NOD SCID mice and lung colonization was determined after 12 days. *Tbk1*-mutant cells were less efficient at forming lung lesions as evidenced by gross lesion formation, by H&E and by lung weight **(Fig. 5a-d)**.

**Figure 5.**
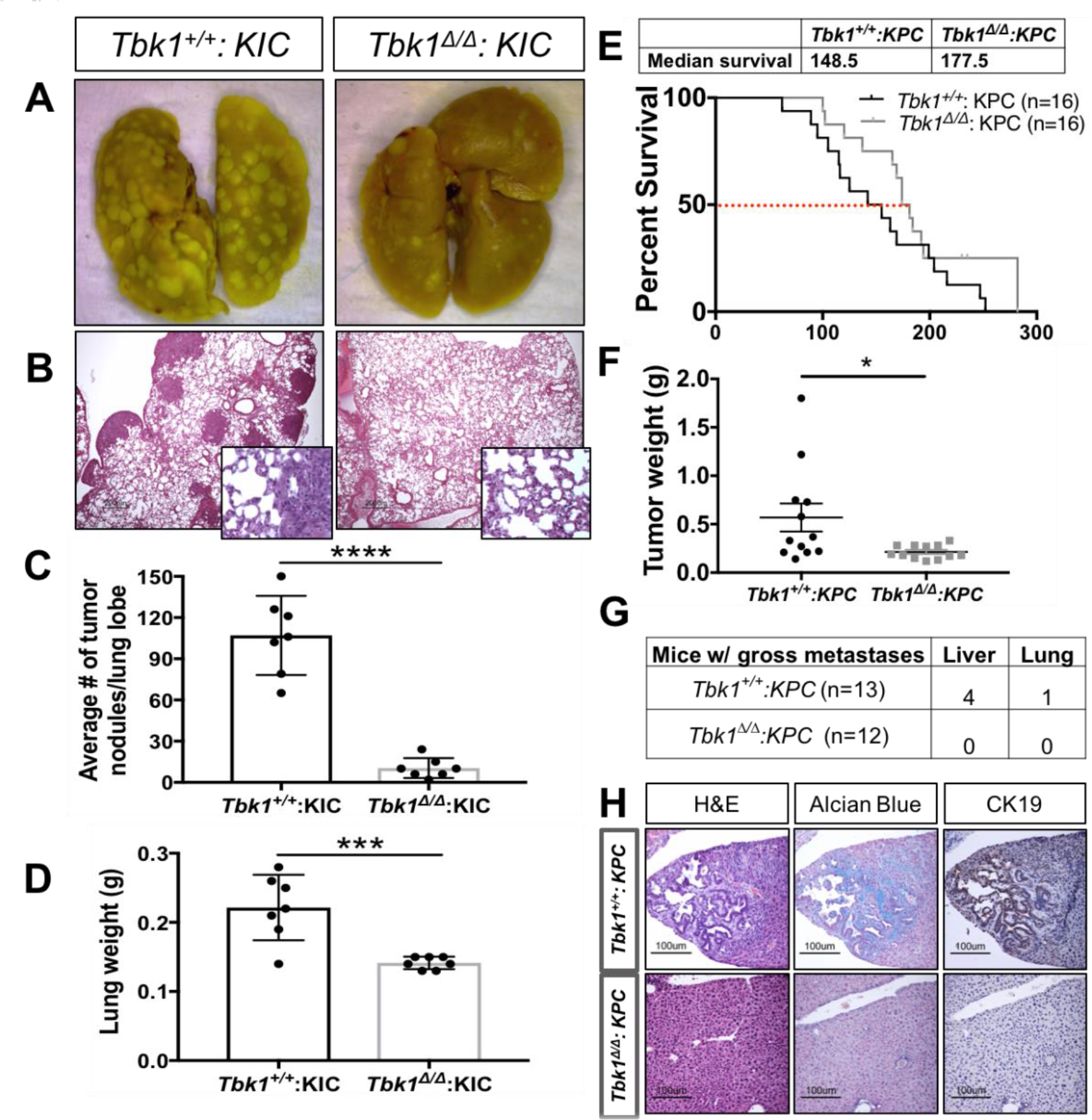
*Tbk1* loss in pancreatic ductal adenocarcinoma (PDA) shows a reduction in metastatic events. **a)** Representative images of lungs from NOD SCID mice sacrificed 12 days after i.v. injection with *Tbk1^+/+^ or Tbk1^Δ/Δ^: KIC* tumor cells (100,000 cells injected per mouse, n = 7 mice/group). **b)** H&E section of lungs from **(a)**, highlighting the tumor nodules. **c)** Number of tumor nodules per lung lobe from **(a)**. **d)** Lung weights from **(a)**. **e)** Kaplan-Meier survival curve of *Tbk1^Δ/Δ^*: *KPC* and *Tbk1^+/+^*: *KPC* mice (n = 16 mice/group). A log-rank Mantel-Cox test was used for survival comparison, *p* = 0.15. **f)** Tumor weights of *Tbk1^Δ/Δ^*: *KPC* and *Tbk1^+/+^*: *KPC* mice. **g)** Number of mice from **(f)** with gross metastases in indicated organs. **h)** Representative liver histology in mice from **(f)**, including H&E, alcian blue, and CK19 immunohistochemical staining. Scale bar indicates 100 μm. Results are representative of mean +/- SEM. Unpaired *t* test, \**p* < 0.05, \*\*\**p* < 0.001, \*\*\*\**p* < 0.0001.

While the experimental metastasis assay results were striking, the effect of *Tbk1* loss on spontaneous metastatic development was also of interest. Therefore, we crossed *Tbk1^Δ/Δ^* mice to *KPC* (*Kras^LSL-G12D/+^*; *LSL-Trp53^LSL-R172H/+^*; *Ptf1a^Cre/+^*) animals. The *KPC* model differs from the *KIC* model in that it contains a dominant negative p53 point mutation instead of loss of the tumor suppressor, *Cdkn2a* [15]. *KPC* mice have a longer median survival (5 months) allowing more time for tumor cells to metastasize [15, 31]. Although not statistically significant, *Tbk1^Δ/Δ^*: *KPC* mice live 1 month longer than *Tbk1^+/+^*: *KPC* mice, shifting the median survival from 5 months to 6 months (*p* = 0.15) **(Fig. 5e)**. Primary tumor burden was significantly reduced in *Tbk1^Δ/Δ^*: *KPC* animals relative to *Tbk1^+/+^*: *KPC* animals **(Fig. 5f)**. Liver and lung metastases were evaluated grossly and by H&E, alcian blue, and CK19 immunohistochemical staining **(Fig. 5g-h)**. As expected, ∼40% of *Tbk1^+/+^*: *KPC* mice were positive for metastasis, but strikingly, no metastatic lesions were detected in *Tbk1^Δ/Δ^*: *KPC* mice **(Fig. 5g-h)**. Consistent with the *KIC* model and the experimental metastasis model, loss of functional *Tbk1* in the *KPC* GEMM restricts tumor cell metastases.

### Re-expression of *Tbk1* in *Tbk1^Δ/Δ^: KIC* cells

To confirm that TBK1 promotes pancreatic tumor cell motility, we stably re-expressed full-length human *TBK1* by lentiviral infection in *Tbk1^Δ/Δ^: KIC* tumor cells and assayed them for invasive and migratory activity. Re-expression of *TBK1* in *Tbk1^Δ/Δ^: KIC* tumor cells was confirmed by Western blot **(Fig. 6a)**. Though the level of *TBK1* re-expression in *Tbk1^Δ/Δ^: KIC* tumor cell lines was substantially lower than endogenous TBK1 levels in *Tbk1^+/+^*: *KIC* cell lines, we did detect a rescue of the mesenchymal morphology in *Tbk1^Δ/Δ^: KIC* cells infected with TBK1-expressing lentivirus (pCDH-TBK1) compared to empty vector-expressing (pCDH-empty vector) cells **(Fig. 6b, Supplemental Fig. 3)**. Further, *Tbk1^Δ/Δ^: KIC* cell lines rescued with pCDH-TBK1 formed 2-3x as many lung tumor nodules, resulting in greater diseased lung burden than pCDH-empty vector-infected cells after i.v. injection **(Fig. 6c-f)**. These results demonstrate that *Tbk1* loss is responsible for the migratory and invasive deficiency in *Tbk1^Δ/Δ^*: *KIC* cells and highlight a novel function for TBK1 in promoting a migratory program in tumor cells.

**Figure 6.**
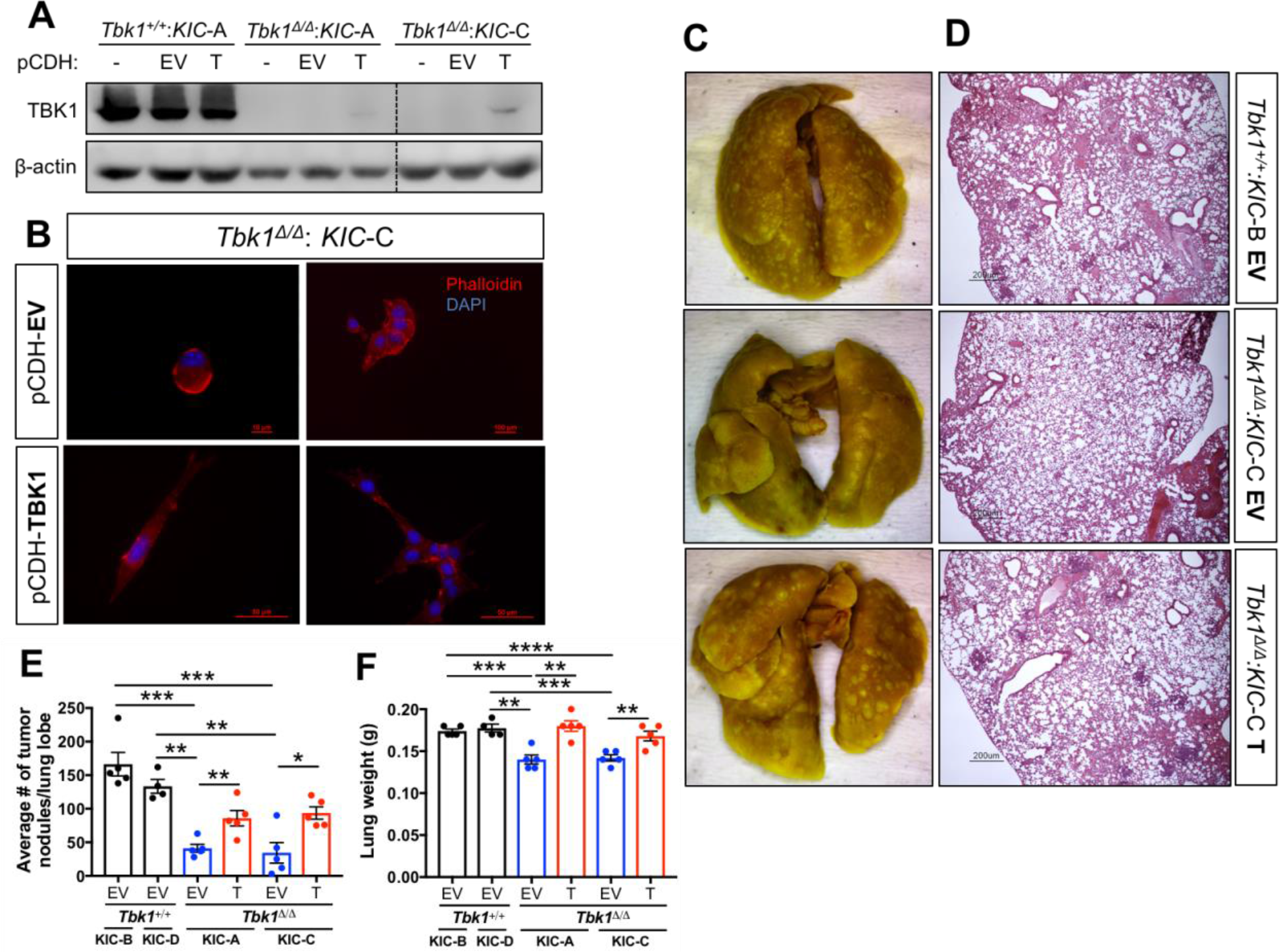
Re-expression of *Tbk1* partially reverses migratory deficit in *Tbk1^Δ/Δ^*: *KIC* cells. **a)** Protein lysates from *Tbk1^Δ/Δ^: KIC* and *Tbk1^+/+^: KIC* cell lines infected with pCDH-empty vector (EV) or pCDH-TBK1 (T) were immunoblotted for TBK1. β-actin was used as a loading control. Dotted line indicates where blot was cropped; however, all samples were run on the same gel and exposed simultaneously. **b)** Representative images of *Tbk1^Δ/Δ^: KIC-C* cells infected with EV or T plated on a mixed layer of collagen and Matrigel. Nuclei are labeled with DAPI (blue) and F-actin is labeled with phalloidin (red). **c)** Representative images of lungs from NOD SCID mice sacrificed 7 days after i.v. injection with *Tbk1: KIC* EV and T-cell lines (100,000 cells injected per mouse, n = 4-5 mice/group). **d)** H&E section of lungs from **(c)** highlighting tumor nodules. Scale bars indicate 200 μm. **e)** Number of tumor nodules per lung lobe from **(c)**. **f)** Lung weights from **(c)**.

### TBK1 is central to Axl-driven epithelial plasticity

Pancreatic tumor cells frequently exploit EMT programs during metastatic dissemination [30, 32]. However, the absence of functional TBK1 in pancreatic tumor cells limits EMT, invasion, and metastases. We recently reported that TBK1 is downstream of the receptor tyrosine kinase Axl, a receptor associated with EMT in PDA [18, 33]. Pharmacological inhibition of Axl led to a concentration-dependent decrease of TBK1 activity while the stimulation of Axl with its ligand, Gas6, resulted in TBK1 activation [18]. To determine if TBK1 is central to Axl-driven EMT, we evaluated Axl signaling in *Tbk1^Δ/Δ^: KIC* tumor cells. Axl was stimulated in *Tbk1^Δ/Δ^: KIC* and *Tbk1^+/+^: KIC* cells with AF854, an activating anti-Axl antibody [22], and the resulting cell lysates were probed for epithelial (E-cadherin, claudin-1), mesenchymal (N-cadherin, Slug) and Axl signaling targets (AKT). Axl activation induced N-Cadherin and Slug protein by 2–3-fold in *Tbk1^+/+^: KIC* tumor cells while having no effect on mesenchymal markers in *Tbk1^Δ/Δ^: KIC* tumor cells **(Fig. 7a)**. Furthermore, pAKT levels increased 5-fold in *Tbk1^+/+^: KIC* cells and remained unaltered in *Tbk1^Δ/Δ^: KIC* tumor cells upon AF854 treatment, indicating that TBK1 may be upstream of AKT. Next, we investigated Axl-induced RAS activation in *Tbk1^+/+^: KIC* cell lines as a possible link between the Axl and TBK1 signaling cascade. *Tbk1^+/+^: KIC-A* cells treated with AF854 showed a substantial increase of GTP-bound RAS, demonstrating that RAS activity is augmented by Axl activation **(Fig. 7b)**. Previous work has shown a RalB GTPase-mediated activation of TBK1 [11]; therefore, we tested if Axl-induced activation of RAS could increase GTP-bound RalB, mediating the activation of TBK1. *Tbk1^+/+^: KIC-A* cells treated with AF854 showed a 3-fold increase in RalB-GTP, demonstrating that RalB activity is increased upon Axl activation **(Fig. 7c)**. Axl-induced Ras and RalB activation was validated in an additional murine PDA cell line, KPfC-8, derived from a spontaneous tumor in the *KPfC (Kras^LSL-G12D/+^*; *Trp53^lox/lox^; Ptf1a^Cre/+^*) GEMM of PDA **(Fig. 7d)**. To extend and confirm these findings in human pancreatic cancer, we stimulated Panc-1 cells with Gas6, which we previously demonstrated increases TBK1 activity [18], and detected an increase in GTP-bound Ras and RalB **(Fig. 7e)**. These results are the first to show that Axl activates RAS and RalB, which leads to downstream activation of TBK1 and AKT, ultimately resulting in increased EMT and a more aggressive and metastatic tumor cell phenotype **(Fig. 7f)**.

**Figure 7.**
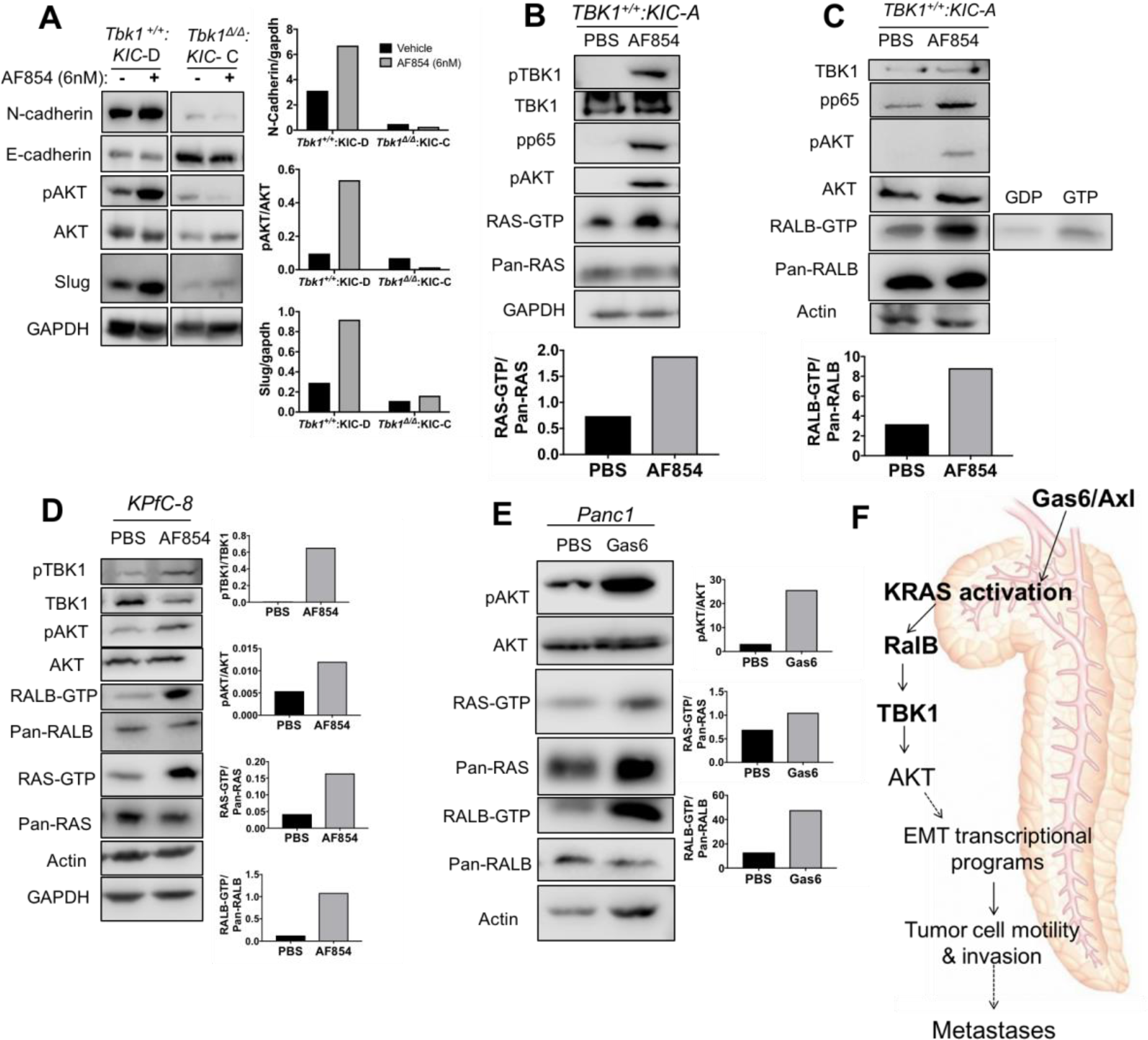
TBK1 promotes epithelial-to-mesenchymal transition (EMT) downstream of Axl. **a)** Protein lysates from *Tbk1: KIC* cell lines treated with PBS or AF854 (6 nM) for 30 min were immunoblotted for indicated proteins and quantified. Protein lysates from *Tbk1: KIC-A* cells treated with PBS or AF854 (6 nM) for 30 min were assayed for active Ras **(b)**, active RalB **(c)**, and immunoblotted for indicated proteins and quantified. GDP and GTP protein loading were used as positive and negative controls, respectively. **d)** Protein lysates isolated from *KPfC-8* cells treated with PBS or AF854 (6 nM) for 30 min were assayed for active Ras, active RalB, and immunoblotted for indicated proteins and quantified. **e)** Protein lysates from Panc-1 cells treated with PBS or Gas6 for 30 min were assayed for active Ras, active RalB, and immunoblotted for indicated proteins and quantified. GAPDH and actin were used as loading controls. **f)** Axl activation in pancreatic cancer cells increases GTP-bound Ras and RalB and subsequent TBK1 and AKT activation, leading to enhanced EMT and ultimately metastatic progression.

## Discussion

Activating mutations in *KRAS* are the dominant oncogenic drivers of pancreatic cancer [3, 4]. No other common epithelial cancer has a single gene with comparable mutation frequency, yet efforts to therapeutically target mutant RAS proteins have not been successful [5]. However, targeting signaling components downstream of RAS that are required for RAS-mediated oncogenesis presents a viable therapeutic alternative [34]. TBK1 is a crucial effector of mutant active KRAS that we found to be expressed abundantly in *KRAS*-mutant PDA tumors and cell lines. TBK1 expression correlated negatively with survival in human pancreatic cancer patients. In addition, the pharmacological inhibition of TBK1 with the selective inhibitor compound II was found to limit human and murine pancreatic tumor cell growth in a mutant *KRAS*-selective manner. The assessment of *Tbk1* loss in multiple clinically relevant GEMMs of PDA revealed that PDA mice lacking kinase active TBK1 have significantly smaller and more epithelial tumors, resulting in fewer metastatic lesions relative to PDA mice with wild-type *Tbk1.* Mechanistic studies established that TBK1 promotes EMT downstream of Axl in PDA, providing insight into a novel function for TBK1. Further, these studies suggest that therapies targeting TBK1 could be used to exploit *KRAS*-mutant tumors.

It is estimated that nearly 90% of cancer mortalities are due to metastases [35], yet TBK1 studies to date have evaluated the effect of TBK1 inhibition only on primary tumor burden. EMT is a hallmark of metastasis in pancreatic cancer and is critical to cancer cell dissemination [36-38]. Within this morphological cellular program, epithelial cancer cells lose contact with the basement membrane and neighboring cells while gaining a more mesenchymal and invasive phenotype [36, 39]. Our results show that tumors and isogenic cell lines from the *KIC* pancreatic GEMM that lack functional TBK1 are more epithelial in gene expression and morphology than *KIC* tumors containing wild-type *Tbk1.* These findings, in combination with mechanistic studies demonstrating that TBK1 is downstream of the EMT driver Axl, indicate that EMT in pancreatic tumor cells is halted by *Tbk1* loss. TBK1 has been linked to EMT in other cancer types. In contrast to our results, knockdown of TBK1 in ERα-positive breast cancer cells reportedly induced EMT and enhanced tumor growth and lung metastasis by suppressing ERα expression [23]. However, in two separate recent studies, gene expression analysis revealed that a mesenchymal gene signature in melanoma and non-small cell lung cancer (NSCLC) cell lines was associated with sensitivity to TBK1 inhibition (TBK1i) [40, 41]. Further analysis revealed mutations in *RAS* family members as a common feature of NSCLC cell lines that showed sensitivity to TBK1i while NSCLC cells that were resistant to TBK1i had a more epithelial gene expression profile and less frequent activating RAS mutations [40]. The mesenchymal gene signature in TBK1i-sensitive NSCLC lines is consistent with our observations in KRAS-driven *Tbk1^+/+^: KIC* tumors that have undergone EMT. Moreover, the epithelial gene expression profile of TBK1-resistant NSCLC cells lines matches the epithelial phenotype of *Tbk1^Δ/Δ^: KIC* tumors that grew independent of TBK1. Though the precise mechanism of how TBK1 promotes EMT is unclear, TBK1 can directly activate AKT [9]. AKT activation can drive EMT via the induction of Snail and Slug that transcriptionally repress E-cadherin and induces Vimentin, Twist1, MMP-2, and MMP-9 that promote tumor cell invasion [39, 42, 43]. Current studies are focused on understanding the interaction between TBK1 and AKT driving the mesenchymal phenotype in PDA and the identification of additional TBK1 substrates that promote EMT programs.

Interestingly, *Tbk1*-mutant PDA tumors were smaller than *Tbk1* wild-type tumors, indicating that *Tbk1* loss affects primary tumor growth in addition to tumor cell motility. TBK1 is central to numerous biological processes that could affect the growth of the primary tumor, including cell division, autophagy, innate immune response, and AKT/mTOR signaling [26, 40, 41, 44-49]. In the context of pancreatic cancer, TBK1 has been reported to promote basal levels of autophagy as a means of silencing cytokine production [48]. These findings imply that the inhibition or loss of TBK1 in PDA could increase cytokine production, ultimately driving immune activation and potentially an antitumor immune response. While we did not investigate the immune landscape of the PDA GEMMs, we did find via gene expression analysis that *Tbk1^Δ/Δ^: KIC* tumors displayed a higher expression of a number of pro-inflammatory genes relative to *Tbk1^+/+^: KIC* tumors **(Supplemental Fig. 4)**. Pro-inflammatory genes with increased expression in *Tbk1^Δ/Δ^: KIC* tumors included *Cxcl1, Ccl2, Ccl4, Ccl27, Irf1*, and *Il1b*. The elevated pro-inflammatory gene expression in *Tbk1^Δ/Δ^: KIC* tumors could be indicative of a heighted inflammatory state and/or an antitumor immune response to some degree. While these results are not sufficient to conclude that *Tbk1* loss promotes antitumor immunity in PDA, the idea is not unreasonable given that *Tbk1^Δ/Δ^* mice have been shown to produce higher levels of pro-inflammatory cytokines in response to immune challenge [12]. In agreement with the notion that *Tbk1* loss produces antitumor immunity, two recent studies reported that immune evasion and metastatic behavior are highly associated with the engagement of the cGAS-STING-TBK1 innate immune pathway in cancer cells [50-52]. In the first study, Backhoum et al. [51] found that chromosomal instability in cancer cells, caused by errors in chromosomal segregation during mitosis, promoted cellular invasion and metastasis through the introduction of double-stranded DNA into the cytosol, engaging the cGAS-STING-TBK1 antiviral pathway. In the second report, Cañadas et al. [52] characterized an interferon-stimulated positive feedback loop of antisense endogenous retroviruses (ERVs) present in a number of human cancer cell lines that produced hyperactive innate immune signaling, myeloid cell infiltration, and immune checkpoint activation. Additionally, they discovered that high ERV-expressing cancer cells correlate with an Axl-positive mesenchymal state, which is consistent with our observations [52].

An important consideration with our *Tbk1^Δ/Δ^* PDA models is that the global *Tbk1* mutation eliminates TBK1 kinase activity and significantly reduces *Tbk1* expression in all cell types, including immune cells, which could impact immune responses to tumor challenge. In fact, a recent study demonstrated that dendritic cell conditional *Tbk1* knockout mice (*Tbk1-DKO*) injected subcutaneously with B16 melanoma cells lived longer and had smaller tumors compared to wild-type *Tbk1* control mice [46]. An assessment of B16 melanoma tumors from *Tbk1-DKO* animals revealed enhanced interferon-responsive gene expression and greater T-effector cell infiltration into tumors and lymph nodes, confirming antitumor immunity conferred by dendritic cell *Tbk1* loss. Collectively, these observations support a pro-tumor immune function for TBK1 that could contribute to the larger tumor sizes in *Tbk1* wild-type PDA mice.

Going forward, it will be important to understand the unique function of TBK1 in each relevant cell type within a tumor. Here we provide evidence of the effects of global *Tbk1* loss in all cell types with our *Tbk1^Δ/Δ^* PDA models, which in many respects is biologically analogous to the effects of pharmacologically inhibiting TBK1 systemically. Overall, our findings expand the spectrum of biological activities of TBK1 and suggest that the therapeutic inhibition of TBK1 may be a useful strategy to control tumor cell invasion and resulting metastases in RAS-driven cancers.

## Author contributions

VHC, ENA, and RAB conceived and designed the study. VHC, ENA, WD, and AEB acquired data and performed analysis and interpretation of data. VHC and ENA wrote the manuscript. RAB reviewed and revised the manuscript. RAB supervised the study.

## Acknowledgements

We thank Drs. Jonathan Cooper and Aubishek Zaman for technical advice and shared resources, Drs. Peiqing Shi and James Chen for the TBK1 expression construct and Dave Primm for editorial assistance. We also thank Dr. Tae Hyun Hwang for assistance with clinical analysis. We gratefully acknowledge Jason Toombs, Tara Billman, Dan Li, and Melissa Gross for their generous assistance with the mouse studies. The work was supported by NIH grants R01 CA192381 and U54 CA210181 Project 2 to RAB, T32 CA124334 (PI: J. Shay) to VHC and support from the Effie Marie Cain Scholarship in Angiogenesis Research and the Gillson Longenbaugh Foundation to RAB. The funders had no role in study design, data collection and analysis, decision to publish, or preparation of the manuscript.

